# Natural SARS-CoV-2 infection in kept ferrets, Spain

**DOI:** 10.1101/2021.01.14.426652

**Authors:** Christian Gortázar, Sandra Barroso-Arévalo, Elisa Ferreras-Colino, Julio Isla, Gabriela de la Fuente, Belén Rivera, Lucas Domínguez, José de la Fuente, José M. Sánchez-Vizcaíno

## Abstract

We found SARS-CoV-2 RNA in 6 of 71 ferrets (8.4%) and isolated the virus from one rectal swab. Natural SARS-CoV-2 infection does occur in kept ferrets, at least under circumstances of high viral circulation in the human population. However, small ferret collections are probably unable to maintain prolonged virus circulation.

## Text

Natural infection of animals with severe acute respiratory syndrome coronavirus 2 (SARS-CoV-2) has been reported in pet cats and dogs, zoo felids, and mustelids belonging to the subfamily mustelinae [*1*]. Among mustelids, natural SARS-CoV-2 infections have been recorded in farmed American mink (*Neovison vison*), and sporadically in a wild mink sampled close to an infected farm in Utah^1^ and in a kept pet ferret (*Mustela putorius furo*) from an infected household in Slovenia^2^. Ferrets are common laboratory models and experimental infections have evidenced their susceptibility and ability to transmit the virus to other ferrets. SARS-CoV-2 is shed up to 8 days post-infection (dpi) in nasal washes, saliva, urine, and feces and is effectively transmitted to naive ferrets by direct contact and via the air [*2, 3*]. Experimentally infected ferrets display either no clinical signs or exhibit elevated body temperature and loss of appetite [*2, 4*].

Ferrets are common pets^3,4,5^, and are also used as work animals for rabbit control. However, it remains unknown if SARS-CoV-2 circulates among kept ferret populations and if ferrets, like farmed mink, could contribute to virus maintenance.

## The Study

We studied 71 ferrets belonging to seven owners and used as working animals for rabbit hunting in Ciudad Real province, central Spain. Group sizes ranged from four to 21 (mean 10). Twenty ferrets belonging to groups 1 and 2 were re-sampled 66 days after initial sampling. Sampling took place between August and November 2020. Animal sampling procedures had been approved by the Madrid Animal Research Ethics Committee, ref. CM14/2020. One oropharyngeal and one rectal swab (DeltaSwab® Virus 3ml, Deltalab S.L., Rubí, Spain) were taken from each ferret for RNA extraction.

SARS-CoV-2-specific RNA was detected using a RT-qPCR assay. Briefly, RNA was extracted using the KingFisher Flex System (Thermo Fisher, Waltham, MA, USA) according to the manufacturer instructions. Detection of SARS-CoV-2 RNA was performed using the envelope protein (E)-encoding gene and two targets (IP2 and IP4) of the RNA-dependent RNA polymerase gene (RdRp) in an RT-PCR protocol established by the WHO according to the guidelines (https://www.who.int/emergencies/diseases/novel-coronavirus-2019/technical-guidance/laboratory-guidance) [*5, 6*]. Primer sets used are detailed in Table 1. The RT-qPCR was carried out using the SuperScript III Platinum One-Step RT-qPCR Kit (ThermoFisher, Massachusetts, USA), according to the manufacturer’s protocol on a CFX Connect™Real-Time PCR Detection System (BioRad, Berkeley, USA). The positive control for real-time RT-qPCR was an *in vitro* transcribed RNA derived from the strain BetaCoV_Wuhan_WIV04_2019 (EPI_ISL_402124), loaned by the Pasteur Institute (Paris, France). Nuclease-free water was used as negative control. A cycle threshold (Ct) cut-off of 40 cycles was used. A result was considered positive when the sample showed a positive RT-qPCR for at least two of the three analyzed targets.

**Table 1.**
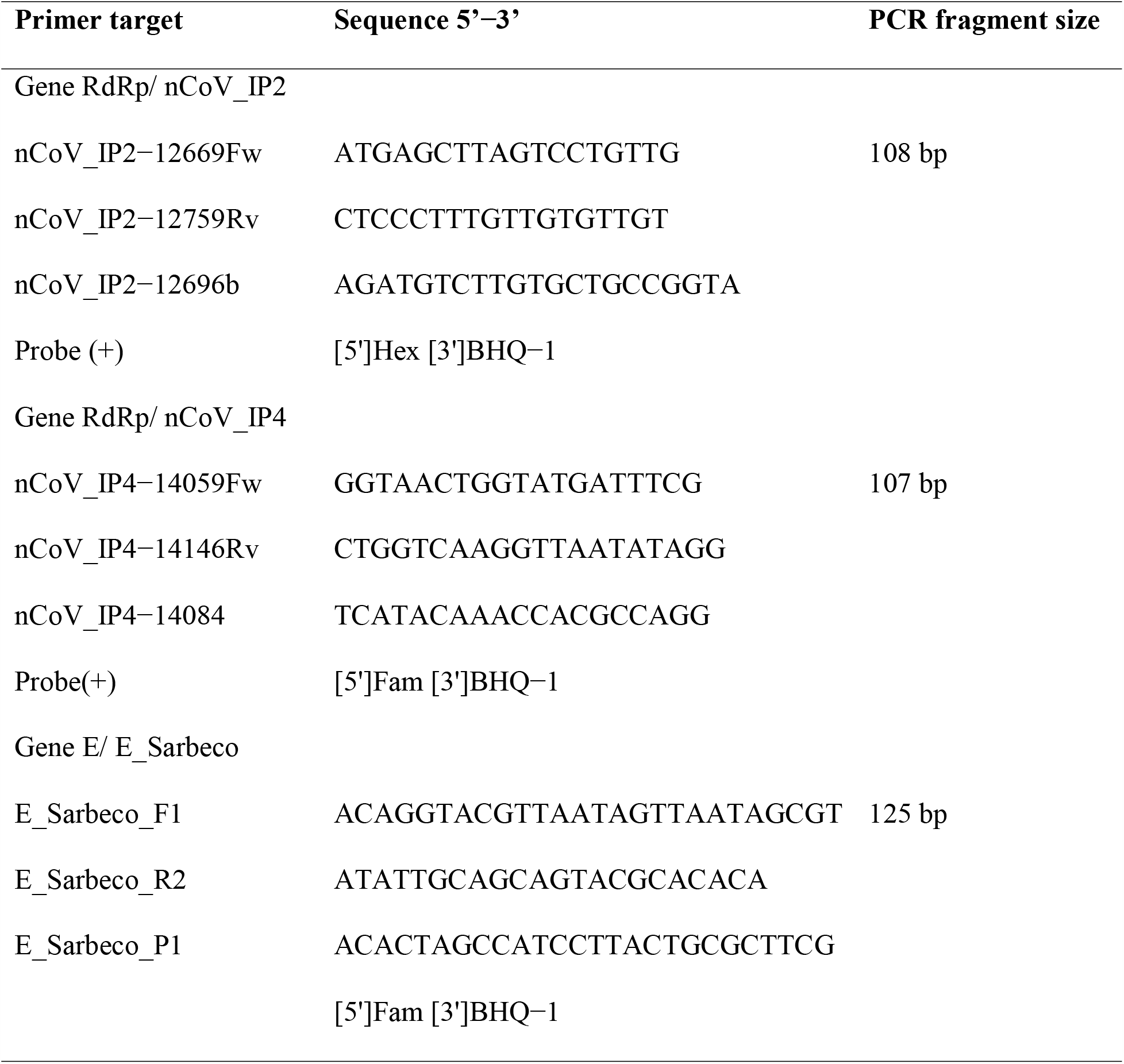
Primer sequences and amplified fragment sizes in base pairs.

Specimens considered positive by RT-qPCR were subjected to virus isolation in Vero E6 cells. Cells were cultured in RPMI 1640 medium supplemented with 10% fetal bovine serum (FBS; Gibco, Madrid, Spain), 100 □IU/ml penicillin, and 100 □μg/ml streptomycin. Cells were seeded in 96-well culture plates and cultured at 37°C with 5% CO_2_ for 24 to 48 h. Then, cells were inoculated with 10 μl of the direct sample (oronasal or fecal swabs). Mock-inoculated cells were used as negative controls. Cultured cells were maintained at 37 °C with 5% CO_2_, with a daily observation of virus-induced cytopathic effect (CPE) and cellular death. After 6 days, cell cultures were frozen, thawed, and subjected to three passages with inoculation of fresh Vero E6 cells with the lysates as described above. SARS-CoV-2 molecular detection was performed by RT-qPCR on the supernatants from every passage to confirm the presence/absence of the virus in the cell culture.

We found SARS-CoV-2 RNA in swab samples from 6 of 71 ferrets (8.4%) (Table 2), belonging to four of seven investigated groups (57%). The likelihood of a swab testing positive was unrelated with age class (under or over one year-old), sex and oral/rectal sample origin (Fisher’s two-tailed p values >0.2). RT-qPCR results were confirmed by sequencing the positive PCR product. None of the 20 re-sampled ferrets was PCR-positive, including one individual that had tested positive two months earlier (oropharyngeal swab; Ct = 35.38).

**Table 2.** RT-qPCR positive results by sample type.

| Animal ID | Sample type | Ct value |
| --- | --- | --- |
| G1-H6 | Rectal swab | 34.5 |
| G1-H17 | Nasal swab | 37.29 |
| G2-H5 | Nasal swab | 35.38 |
| G5-H11 | Nasal swab | 39.83 |
| G7-H7 | Nasal swab | 30.59 |
| G7-H9 | Nasal swab | 38.91 |

SARS-CoV-2 was isolated only from the rectal swab of one ferret (Ct in the original sample = 34.5). Cell culture showed CPE and cellular death in the three passages. Virus recovery was also confirmed by RT-qPCR (Ct value reduction from original inoculum to cell suspension of third passage).

We conclude that natural SARS-CoV-2 infection in kept ferrets does occur in circumstances of high viral circulation in the human population [*7*]. However, the high Ct values observed, and the lack of positive ferrets at re-sampling, indicate that small ferret populations are not as able to maintain prolonged virus circulation as large, farmed mink populations [*8*]. Specific guidance on SARS-CoV-2 in ferrets has been made available in the UK^6^.

## Acknowledgments

This study received funding from ISCIII, Spanish Government, grant number: COV20/01385 and EFC was supported by a grant from Universidad de Castilla-La Mancha, Spain.

## Conflicts of interest

The authors have no conflicts of interest to declare.

## Author Bio

Professor Christian Gortázar heads the SaBio research group at the Spanish Wildlife Research Institute (IREC). His research interests include the epidemiology and control of infections shared between wildlife, livestock, and human beings.

## Footnotes

1 https://promedmail.org/promed-post/?id=8015608.

2 https://www.oie.int/wahis_2/public/wahid.php/Reviewreport/Review?page_refer=MapFullEventReport&reportid=37289.

3 https://www.vettimes.co.uk/app/uploads/wp-post-to-pdf-enhanced-cache/1/overview-of-ferrets-part-one-conditions-and-behaviour.pdf.

4 https://www.avma.org/resources-tools/reports-statistics/us-pet-ownership-statistics.

5 https://www.mapa.gob.es/es/ganaderia/temas/produccion-y-mercados-ganaderos/20160222_informeestudioparapublicar_tcm30-104720.pdf

6 http://apha.defra.gov.uk/documents/guidance-sars-cov-2-ferrets.pdf

## Notes

### Competing Interest Statement

The authors have declared no competing interest.

